# Tissue repair in the mouse liver following acute carbon tetrachloride depends on injury-induced Wnt/β-catenin signaling

**DOI:** 10.1101/507418

**Authors:** Ludan Zhao, Yinhua Jin, Katie Donahue, Margaret Tsui, Matt Fish, Catriona Y. Logan, Bruce Wang, Roel Nusse

## Abstract

In the liver, Wnt/β-catenin signaling is involved in regulating zonation and hepatocyte proliferation during homeostasis. We have examined Wnt gene expression and signaling after injury and we show by *in situ* hybridization that Wnts are activated by acute carbon tetrachloride (CCl_4_) toxicity. Following injury, peri-injury hepatocytes become Wnt-responsive, expressing the Wnt target gene Axin2. Lineage tracing of peri-injury Axin2+ hepatocytes shows that during recovery, the injured parenchyma becomes repopulated and repaired by Axin2+ descendants. Using single cell RNA sequencing (scRNA-seq), we show that endothelial cells are the major source of Wnts following acute CCl_4_ toxicity. Induced loss of β-catenin in peri-injury hepatocytes results in delayed repair and ultimately to injury-induced lethality, while loss of Wnt production from endothelial cells leads to a delay in the proliferative response after injury.

**Conclusion:** Our Pindings highlight the importance of the Wnt/β-catenin signaling pathway in restoring tissue integrity following acute liver toxicity and establishes a role of endothelial cells as an important Wnt-producing regulator of liver tissue repair following localized liver injury.

---

As a major site for both nutrition and xenobiotic metabolism, the liver serves as a first line of defense against environmental toxins. However, the liver can also be easily damaged by the compounds which it metabolizes. As such, the ability to repair damaged liver tissue has garnered much interest. What is the source of new hepatocytes after injury? And how is hepatocyte re-population regulated during injury repair?

In the context of many injury models, mature hepatocytes become activated to generate new hepatocytes during injury repair (1,2). It has also been suggested that following certain forms of liver injury and/or under conditions where hepatocyte proliferation is impaired, bi-potential progenitor cells arising in the periportal region are activated to give rise to both cholangiocytes and hepatocytes, thus serving as a source of new cells in injury repair (3–7). However, the source of these bi-potential cells has been debated (2,6). In either case, the pro-proliferative signals induced by injury to activate hepatocyte proliferation remains to be fully characterized.

The Wnt/β-catenin signaling pathway has been implicated in the regulation of liver biology from normal development to disease (8–11). Previous work, including from our lab, has shown that endothelium-derived Wnt signaling regulates pericentral hepatocyte proliferation during homeostasis. We showed that Wnts secreted by central vein endothelial cells maintain pericentral Axin2+ hepatocytes in a proliferative state (12). In the context of partial hepatectomy, it has been shown that β-catenin is necessary for early post-injury hepatocyte DNA synthesis (13–15). This effect following partial hepatectomy is regulated by macrophage (13) and endothelium-derived Wnts (16), and is further modulated by the Wnt co-activator, Rspo3, which signals through the Lgr5 receptor (17,18). However, the role of Wnt/β-catenin signaling following acute localized liver injury, such as that following carbon tetrachloride toxicity, remains to be fully understood.

Given the importance of the Wnt/β-catenin pathway in regulating hepatocyte proliferation, we asked whether this pathway is involved in regulating proliferation following acute CCl_4_ injury.

## Experimental Procedures

### Animals and carbon tetrachloride injury

All injury experiments were performed on male mice, 8-12 weeks of age. All animals received a single intraperitoneal injection of 1mL/kg CCl_4_ (Sigma, diluted 1 to 4 in corn oil and Piltered using a 0.2μm Pilter prior to administration), unless otherwise stated. Administration of 1mL/kg Piltered corn oil was used for all uninjured controls. All mice were in the C57/BL6 background unless otherwise specified. C57/BL6J wild-type mice from The Jackson Laboratory were used for all experiments unless otherwise specified. Axin2-LacZ (19) and Axin2Cre^ERT2^ (20) mice have been described. Ai9 (B6.Cg-*Gt(ROSA)26Sor^tm9(CAG-tdTomato)Hze^*/J) (21), β-catenin^flox^ (B6.129-*Ctnnb1^tm2Kem^*/KnwJ) (22), Wls^flox^ (129S-*Wls^tm1.1Lan^*/J) (23), and CMV-Cre (B6.C-Tg(CMV-cre)1Cgn/J) (24) mice were obtained from The Jackson Laboratory. Cdh5(PAC)Cre^ERT2^ (25) were obtained from R. Adams. All alleles were heterozygous, except where stated.

To generate mice harboring deletion alleles for β-catenin (β-catenin^Δ^) or Wntless (Wls^Δ^), homozygous β-catenin^flox/flox^ (or Wls^flox/flox^) mice were crossed to CMV-Cre mice. Genotyping PCR to detect the deletion allele was carried out as described (22,23).

For post-injury lineage tracing studies, Axin2Cre^ERT2^; Ai9 mice received a single intraperitoneal injection of tamoxifen (Sigma, 4mg tamoxifen per 25g body weight, dissolved in 10% ethanol/corn oil) 24 hours after CCl_4_ administration, n=3 animals per timepoint.

For β-catenin knock-out studies, CCl_4_ injured Axin2Cre^ERT2^; β-catenin^flox/Δ^ (Bcat-cKO) and β-catenin^flox/Δ^ littermate control mice received daily intraperitoneal 4mg/25g tamoxifen injections for 3 or 4 days, starting on the day of injury. Mice were sacrificed at day 2 (control n=3, Bcat-cKO n=3) or day 4 (control n=3, Bcat-cKO n=5) after injury. For injury survival studies, Bcat-cKO animals that received 1mL/kg corn oil and 4 daily doses of tamoxifen were used for uninjured Bcat-cKO controls.

For Wntless knock-out studies, Cdh5(PAC)Cre^ERT2^; Wls^flox/Δ^ (n=3) and Cdh5(PAC)Cre^ERT2^; Wls^+/flox^ or Wls^+/flox^ littermate control (n=4) mice received a single intraperitoneal injection of 0.75mL/kg CCl_4_ and daily doses of 4mg/25g tamoxifen for 3 days. All animals also received an intraperitoneal dose of 50mg/kg of 5-ethynyl-2’-deoxyuridine (EdU; Invitrogen) three hours prior to sacrifice.

All animal experiments and methods were approved by the Institutional Animal Care and Use Committee at Stanford University in accordance with NIH guidelines. All mice were housed in recyclable individually ventilated cages in Stanford University animal facility on a 12-h light/dark cycle with ad libitum access to water and normal chow. All interventions were done during the light cycle.

### Histology & immunofluorescence

For liver morphology analysis, collected liver samples were Pixed in 10% neutral buffered formalin overnight at room temperature, serially dehydrated in ethanol, cleared in HistoClear (Natural Diagnostics), and paraffin-embedded in Paraplast Plus (Sigma). Paraffin sections (5 μm) were stained with Mayer ‘ s hematoxylin and counterstained with eosin.

For detection of LacZ expression in Axin2-LacZ samples, whole livers were fixed in 1% PFA overnight at 4°C, washed in detergent rinse (PBS with 2 mM MgCl2, 0.01% sodium deoxycholate and 0.02% NP-40) and stained in X-gal staining solution (PBS with 2 mM MgCl2, 0.01% sodium deoxycholate, 0.02% NP-40, 5 mM potassium ferricyanide, 5 mM potassium ferrocyanide and 1 mg/ml 5-bromo-4-chloro-3-indolyl-β-D-galactopyranoside) in the dark at room temperature for 4 days for full tissue penetration. Following staining, tissues were washed, post-fixed in 4% PFA for 1 hour at room temperature and processed for paraffin embedding.

For immunofluorescence antibody staining, samples were fixed in 4% paraformaldehyde overnight at 4°C, cryo-protected in 30% sucrose in PBS for 24 hours at 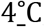, and then embedded in OCT and snap frozen. Cryo-sections (8-10 μm) were blocked in 5% normal donkey serum in 0.5% Triton-X in PBS at room temperature and stained with primary and secondary antibodies, then mounted in Prolong Gold with DAPI mounting medium (Life Technologies). Mouse on Mouse (M.O.M.) detection kit (Vector Labs) was used for all mouse primary antibody staining. The following primary antibodies were used: GS (mouse, 1:500, Millipore MAB302), Glt1 (rabbit, 1:100, Frontier Institute Glt1-Rb-Af670), HNF4a (mouse, 1:100, Abcam ab41898), beta-Actin (rabbit, 1:250, Abcam ab8227), Tbx3 (goat, 1:50, Santa Cruz sc-17871), and Alexa Fluor 488 phalloidin (Thermo Fischer A12379). EdU detection was performed according to the Click-iT EdU Alexa Fluor 647 Imaging Kit (Life Technologies) protocol. All immunofluorescence staining was performed in the dark.

### RNA isolation and qPCR

Liver samples were homogenized in TRIzol (Invitrogen) using a benchtop bead homogenizer and RNA isolation was carried out per product protocol. Isolated RNA was purified using RNeasy Mini Isolation Kit (Qiagen) per product protocol. RNA reverse transcription was performed using High Capacity Reverse Transcription Kit (Life Technologies), and qPCR was performed using duplex Taqman Gene Expression Assays (Axin2, Wnt2, Wnt4, Wnt5a, Wnt9b, Albumin, Ki67, and GAPDH; Life Technologies), run in 96-well plate format on StepOnePlus Real-Time PCR System (Applied Biosystems). No template controls were also performed. Relative target gene expression levels were calculated using the delta-delta CT method (26). The gene expression level of the mutant was normalized to that of the control. Statistical significance in relative expression levels were analyzed with one-way ANOVA in GraphPad Prism 7.

### RNA Scope *in situ*

Paraffin-embedded liver sections were processed for RNA *in situ* detection using either RNAscope 1-plex Detection Kit (Chromogenic) for single *in situ* experiments or RNAScope 2.5 HD Duplex Kit for double *in situ* experiments were used according to manufacturer ‘ s protocol (Advanced Cell Diagnostics). The following RNAscope probes were used: Axin2 (NM 015732, region 330-1287), Wnt2 (NM 023653, region 857-2086), Wnt4 (NM 009523.2, region 2147-3150), Wnt5a (NM 009524.3, region 200-1431), Wnt9b (NM 011719, region 727-1616), Pecam1 (NM 001032378.1, region 915-1827), Adgre1 (NM 010130.4, region 85-1026), Reelin (NM 011261.2, region 9371-10257),_DapB (negative control, EF 191515, region 414-862), and Polr2a (positive control, NM 009089.2, region 2802-3678).

### Single cell RNA sequencing

Single cell suspensions were generated from day 3 CCl_4_-injured mice by a modified two-step collagenase perfusion technique as previously described (12). Hepatocytes were removed by low-speed centrifugation. Non-parenchymal cells were captured as single cells and sequenced with the 10x Genomics droplet-based method (27). CellRanger version 2.1.1 was used to align raw reads and to filter cell barcodes. Quality control, principal component analysis, clustering, and differential expression analysis were performed in R with the Seurat version 2.3.4 package (27). Samples with disproportionally high cell counts were randomly downsampled to the group average. Cells with less than 200 or greather than 3000 genes, UMI count of less than 500 or greater than 10000, and greater than 25% of total expression from mitochondrial genes were filtered out. An initial round of clustering was performed to identify and remove hepatocytes. The remaining cells were sub-clustered and annotated using markers for known non-parenchymal cell types in the liver (Supplemental Figure 5D).

### Microscopy and imaging

All immunohistochemistry, RNA in situ, and immunostaining sections were imaged using a Zeiss Axioplan 2 microscope equipped with AxioCam MRm (fluorescence) and MRC5 (bright field) cameras. Image acquisition was done using Axiovision AC software (Release 4.8, Carl Zeiss).

### Image quantification

For lineage tracing studies, thresholding was applied to single-channel fluorescent images and the pixel area of the thresholded region was measured using ImageJ. “Percent labeled area” was calculated as percent total tdTomato-positive pixel area over total intact tissue area. For day 3 samples, total intact tissue area was defined as total tissue area minus necrotic tissue area. Ten or more representative images were analyzed per animal per timepoint.

For Bcat-cKO tissue repair analysis, H&E images were used for quantification. “Percent lobular repair” was defined as 1-[(injury distance) / (hepatic lobule distance)] x 100%. Hepatic lobule distance was measured as distance from central vein to portal vein. Injury distance was measured as distance from central vein to the injury border. A minimum of 10 hepatic lobules were analyzed per animal per timepoint.

Statistical analysis was conducted using a two-tailed t-test on GraphPad Prism 7. All graphs were generated in GraphPad Prism 7.

## Results

### Injury induces *de novo* Wnt response in mid-lobular hepatocytes

Previously, we had found that in the uninjured liver, pericentral Wnt-regulated Axin2+ hepatocytes proliferate at a higher rate compared to other cells (12). In our current studies, we asked whether Wnt/β-catenin signaling performs a similar role in regulating hepatocyte proliferation following acute liver injury. CCl_4_ injury leads to centrilobular necrosis within 48 hours of CCl_4_ administration (Supplemental Figure 1). The local effect of CCl_4_ is due to the zonated expression of CYP2E1, the major P450 cytochrome that metabolizes CCl_4_ (28). In order to determine whether Wnt signaling is activated following acute CCl_4_ injury, we conducted *in situ* hybridization for the Wnt target gene, Axin2 (19,29). While expression of Axin2 is normally restricted to pericentral hepatocytes (Figure 1A), its expression became undetectable as early as 6 hours following CCl_4_ administration (Figure 1B). However, Axin2 expression re-appears in the injured liver by 3 days following injury, but its expression is observed in *mid-lobular* hepatocytes as opposed to pericentral hepatocytes (Figure 1C). Additionally, these injury-induced Axin2+ hepatocytes encircle the local necrotic tissue, extending from directly adjacent to the injury border to a few cell diameters away (Figure 1C, arrows). This post-injury upregulation of Axin2 expression is consistent with whole liver qRT-PCR results which show significant upregulation of Axin2 expression at 3 days after injury (Figure 1D). Injury-induced expression of Axin2 is sustained throughout the injury repair time course (Figure 1D), indicating that Wnt signaling is active during the entirety of injury repair.

**Figure 1.**
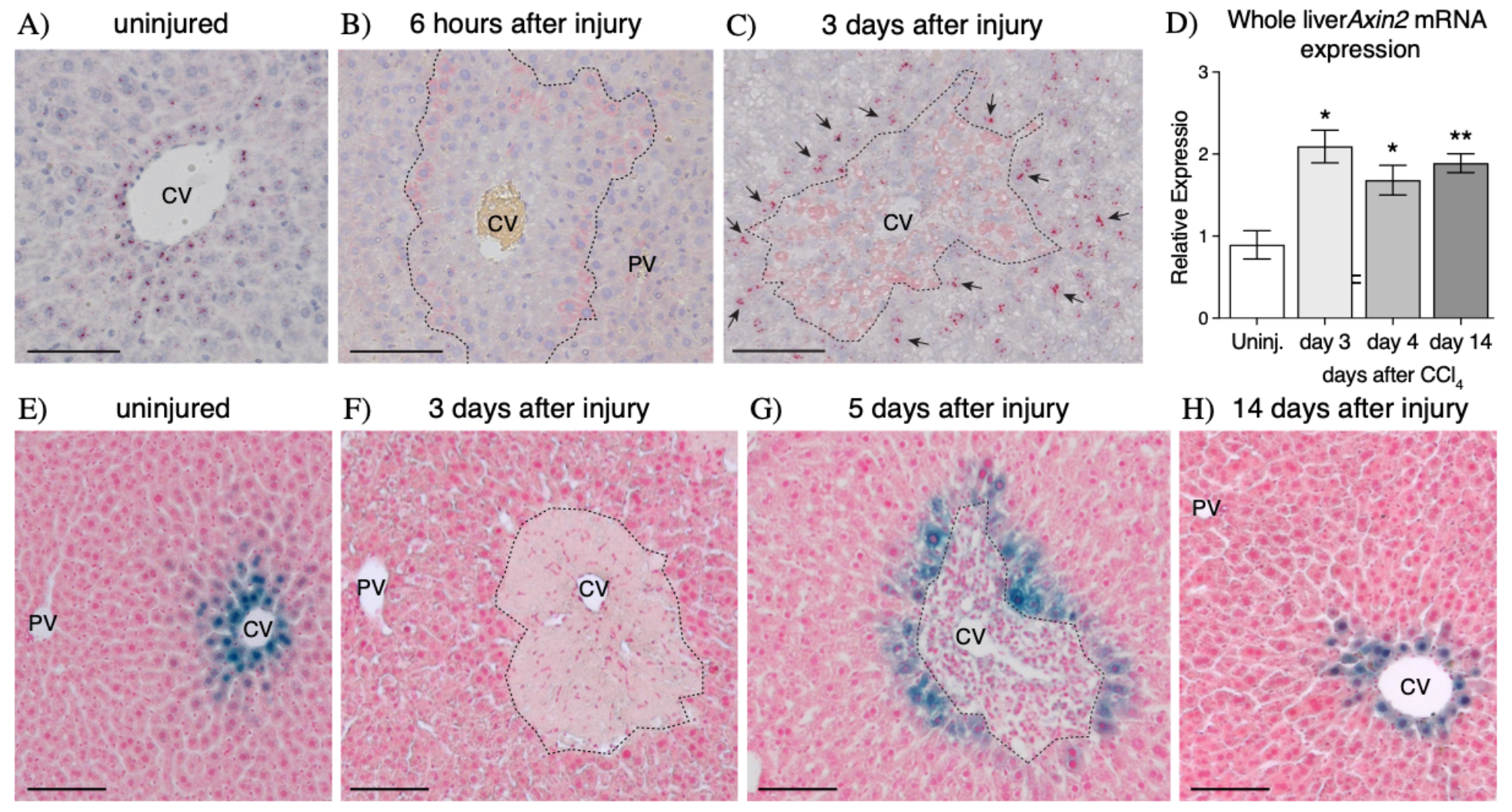
CCl_4_ injury induces *de novo* Axin2 expression in peri-injury hepatocytes. (A-C) Axin2 mRNA *in situ* hybridization in uninjured (A), early injured, and repairing livers show that Axin2 expression is lost as soon as 6 hours after injury (B) but is re-activated in peri-injury hepatocytes during injury repair (C, arrows). (D) Upregulation of Axin2 expression following injury is confirmed by whole liver qRT-PCR. (E-H) Axin2-LacZ reporter staining also confirms the dynamics of *de novo* Axin2 expression in peri-injury hepatocytes following CCl_4_ toxicity. 100μm scale bars shown. CV = central vein, PV = portal vein. Dashed line represents injury border. Error bars show S.E.M. *p<0.05, **p<0.01 by two-tailed t-test compared to uninjured control for n=3 animals per timepoint.

Consistent with our *in situ* hybridization results, Axin2-LacZ reporter animals showed that while the original population of pericentral Axin2+ hepatocytes (Figure 1E) is lost following CCl_4_ administration (Figure 1F), surviving mid-lobular hepatocytes adjacent to the injury border begin to express Axin2 during injury repair (Figure 1G). Interestingly, this pattern of injury-induced Axin2 expression in hepatocytes bordering the damaged tissue is maintained throughout tissue repair (Supplemental Figure 2). Upon full injury recovery, the normal homeostatic pattern of Axin2 expression, restricted to the peri-central cells, is re-established (Figure 1H). Thus, CCl_4_ injury induces activation of a Wnt signaling response in surviving hepatocytes immediately adjacent to the damaged centrilobular region.

In the uninjured liver, pericentral Axin2+ hepatocytes express both glutamine synthetase (GS) and glutamate transporter 1 (Glt1) (Figure 2A & 2E). Upon pericentral hepatocyte ablation following injury, GS and Glt1 are no longer expressed by the surviving hepatocytes in the liver lobule (Figure 2B & 2F). Over the course of injury repair, while rare peri-injury hepatocytes exhibit expression of GS (Figure 2C, arrowhead), Glt1 expression remains negative (Figure 2G). Finally, upon full injury recovery at 14 days following CCl_4_, the homeostatic GS and Glt1 expression patterns are re-established in pericentral hepatocytes (Figure 2D & 2H). These results provide evidence that injury-induced Axin2+ hepatocytes, which are located in the mid-hepatic lobule-adjacent to the injury border, do not express GS or Glt1, <u>and</u> are thus distinct from pericentral Axin2+ cells of the uninjured liver.

**Figure 2.**
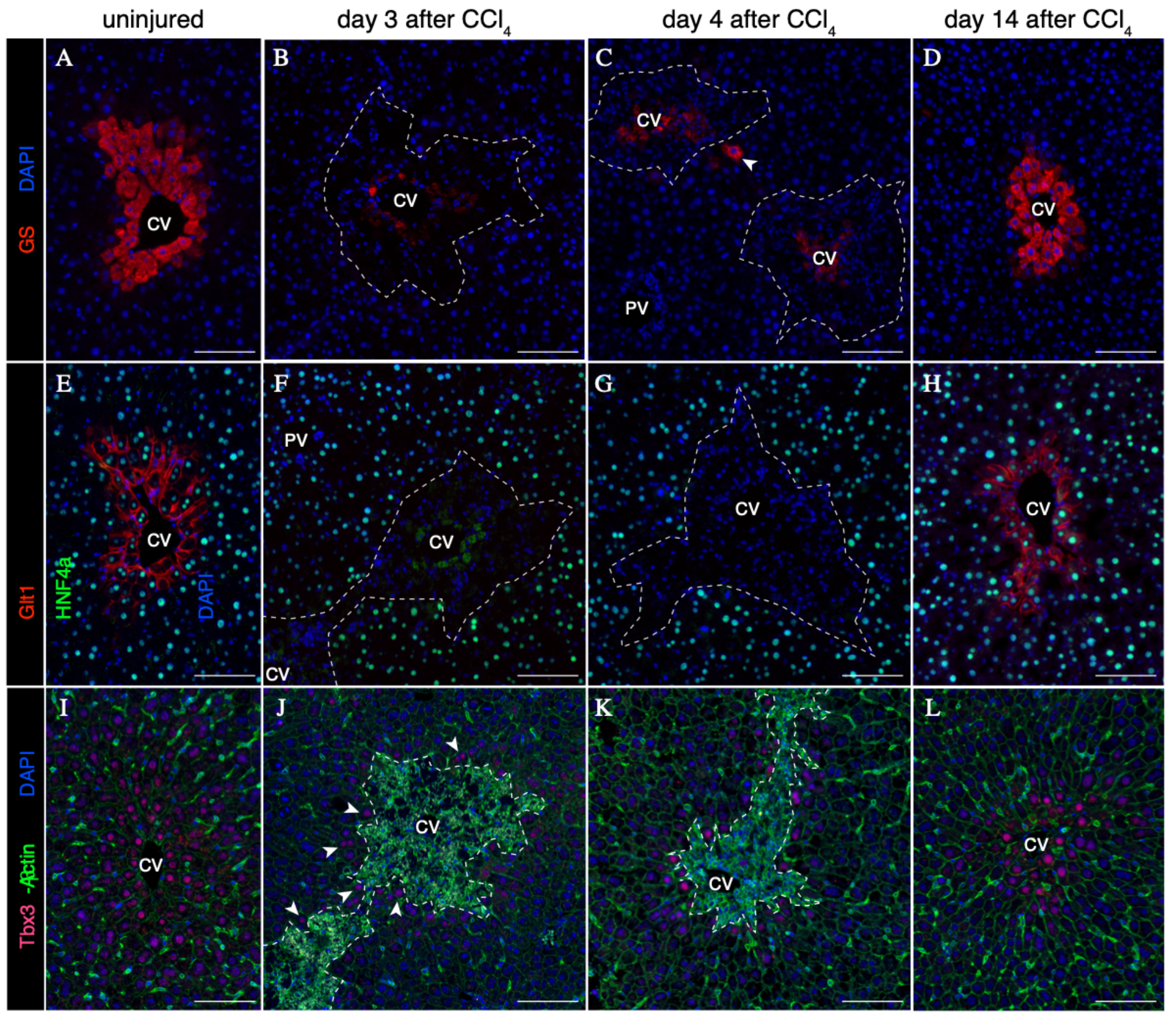
Post-CCl_4_ peri-injury hepatocytes do not express Wnt targets GS and Glt1, but are positive for Tbx3. (A-D) Expression of GS in uninjured (A) and post-injury (B-D) liver. Rare GS+ hepatocytes are observed during injury repair (C, arrowhead). (E-H) Expression of Glt1 in uninjured (E) and post-injury (B-D) liver. (I-L) Expression of Tbx3 in uninjured (I) and post-injury (J-L) liver. Low levels of Tbx3 expression are induced in peri-injury hepatocytes as early as 3 days following injury (J, arrowheads). 100μm scale bars. Dashed line represents injury border. CV = central vein, PV = portal vein. Representative images from n=3 animals per timepoint.

Previously, we reported that pericentral Axin2+ hepatocytes in the uninjured adult liver express the early hepatocyte marker T-box transcription factor 3 (Tbx3) (12) (Figure 2I). Strikingly, while pericentral Tbx3+ hepatocytes are lost following injury, Tbx3 expression is turned on in peri-injury hepatocytes throughout the injury repair process (Figures 2J & 2K). By 14 days following injury, the pericentral Tbx3 expression pattern is re-established (Figure 2L).

### Peri-injury Axin2+ hepatocytes proliferate to repair local injury

To determine whether injury-induced peri-injury Axin2+ hepatocytes proliferate in response to injury, we performed lineage tracing of Axin2+ hepatocytes, using tamoxifen-inducible Axin2Cre^ERT2^; Ai9 reporter mice, after CCl_4_ injury (Figure 3A). At 3 days following injury, peri-injury Axin2+ hepatocytes were labeled with cytoplasmic tdTomato (6.0% ±2.6% of total area, Figure 3B & 3D). Upon full recovery at 14 days after injury, labeled hepatocytes and their progeny accounted for 28.4%±4.2% of the total liver tissue (Figure 3D). Significantly, all central veins at 14 days after injury were lined with rings of labeled hepatocytes (Figure 3C & 3C inset), indicating that peri-injury Axin2+ hepatocytes are the main source of repopulation of the damaged centrilobular tissue.

**Figure 3.**
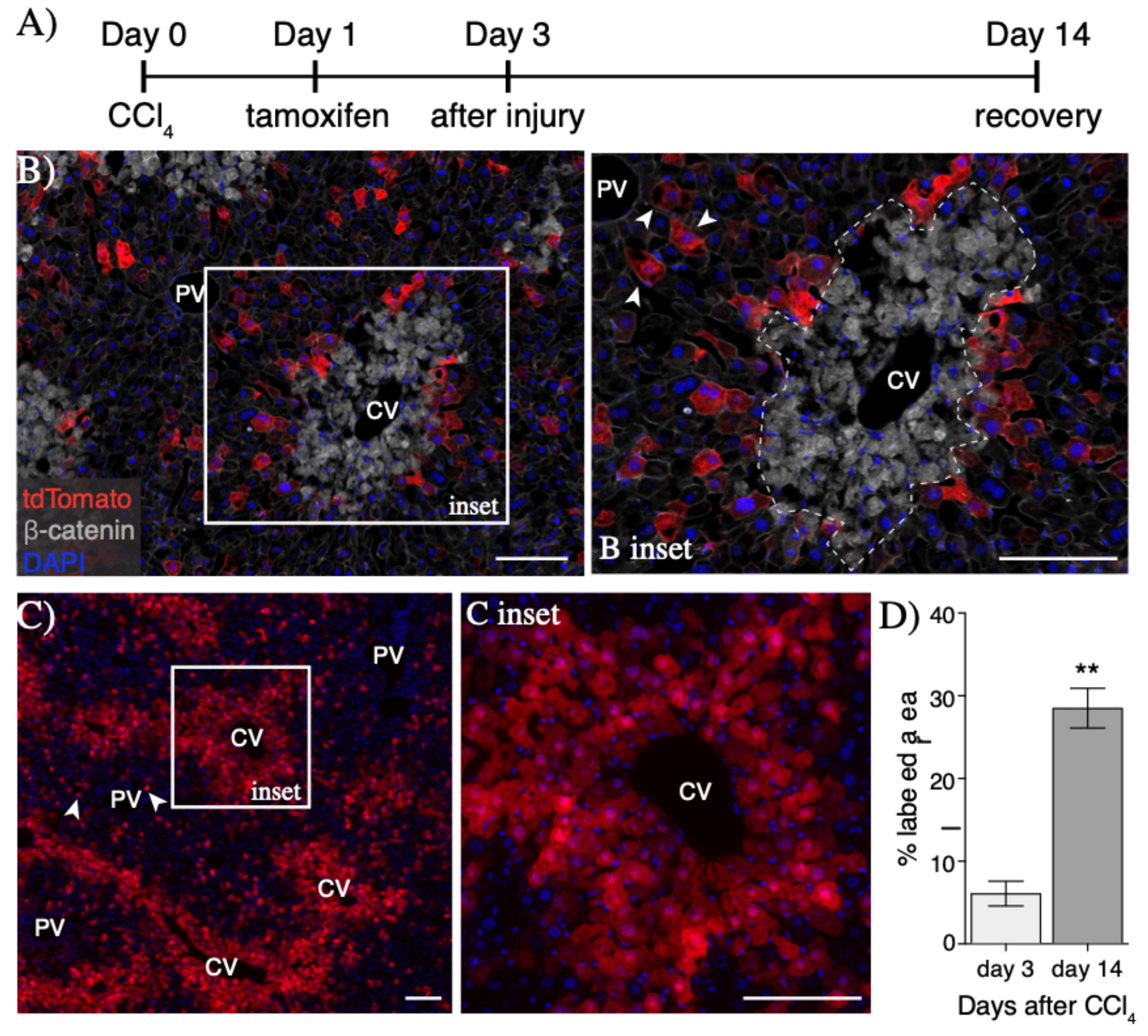
Peri-injury Axin2+ hepatocytes proliferate to repair damaged tissue. (A) Injury and lineage labeling scheme for panels B-D. Peri-injury Axin2+ hepatocytes are efficiently labeled following injury (B) and show significant expansion over the course of tissue repair (C). Notably, tdTomato+ hepatocytes in the periportal space (B inset, arrowheads) show no expansion upon injury repair (C, arrowheads). (D) Quantification of percent labeled area. 100μm scale bars. CV = central vein, PV = portal vein. Dashed line represents injury border. Error bars show S.E.M. **p<0.01 by two-tailed t-test for n=3 animals per timepoint.

We also found that some periportal hepatocytes were labeled following injury (Figure 3B inset, arrowheads). However, these labeled periportal cells did not expand into clones upon injury recovery (Figure 3C, arrowheads). The lack of clone generation by labeled periportal hepatocytes suggest that it is peri-injury hepatocytes, and not hepatocytes elsewhere in the hepatic lobule, that are activated to proliferate following injury.

### Upregulation of endogenous and exogenous Wnt ligands following injury

In the uninjured liver, Wnt2 and Wnt9b are expressed in the endothelial cells lining the central vein (12), while Wnt2 is also expressed in sinusoidal endothelial cells (12,16,30). We conducted a qRT-PCR Wnt screen at various time points following injury to identify those Wnts whose expression may be upregulated following CCl_4_ injury. We found that Wnt2, 4, 5a, and 9b were significantly upregulated following injury and throughout injury repair (Figure 4A). *In situ* hybridization studies on injured liver sections confirmed the expression of these four Wnt ligands within and/or around the necrotic lesion (Figure 4B and Supplemental Figure 4). While the expression of Wnt2 and Wnt4 persisted at higher levels following injury repair 2 weeks after CCl_4_ administration, the expression of Wnt5a and Wnt9b returned to normal levels by 14 days after injury (Figure 4A and Supplemental Figure 4).

**Figure 4.**
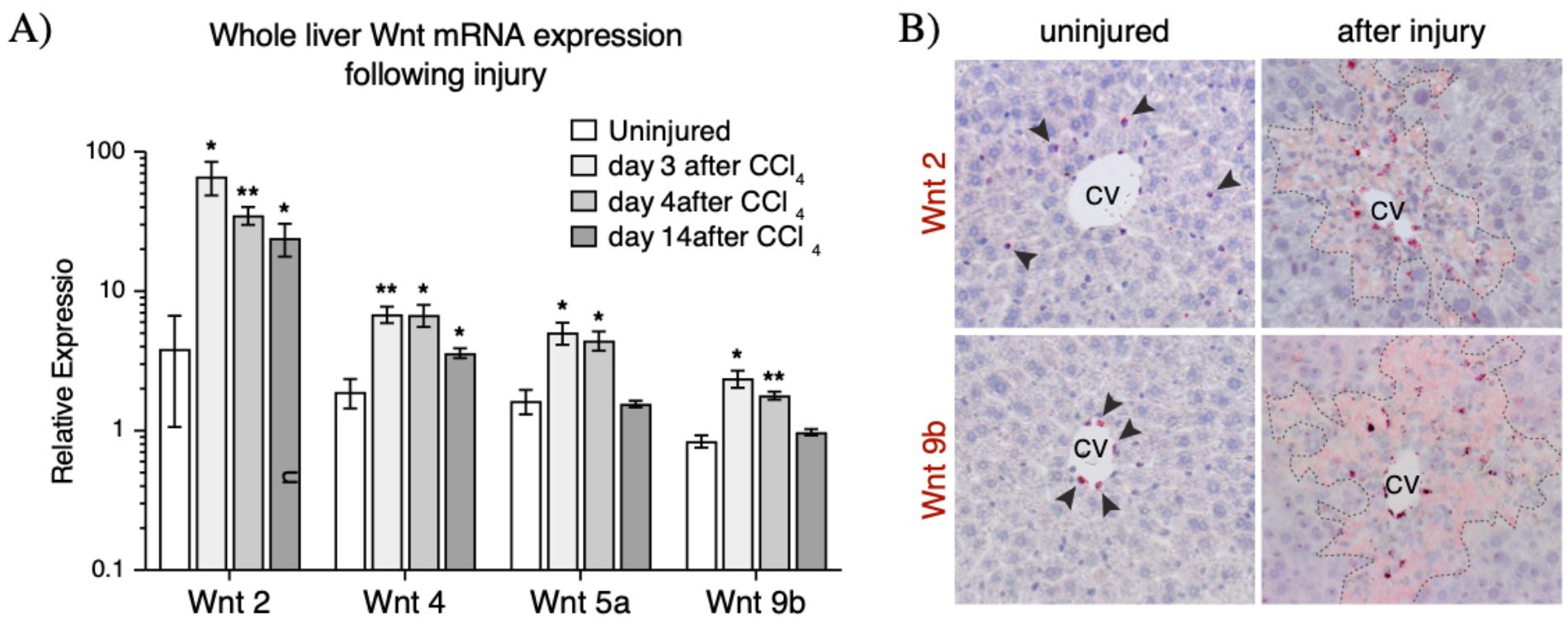
Upregulation and expression pattern of Wnt ligands after injury. (A) Whole liver qRT-PCR results showed significant (p-val<0.05) upregulation of four Wnt ligands whose expression is found in the pericentral and mid-lobular regions following injury. (B) mRNA *in situ* hybridization for Wnt2 and Wnt9b show expanded regions of expression for these two pericentral Wnt ligands early after injury. Arrowheads indicate positive signal in the uninjured liver. *In situ* hybridization images for Wnt4 and Wnt5a can be found in Supplemental Figure 4. Dashed lines represent injury border. Error bars show S.E.M. *p<0.05, **p<0.01 by two-tailed t-test for n=3 animals per timepoint.

The liver-endogenous Wnts, Wnt2 and 9b, were found to be expressed in the locally damaged tissue during both the early necrotic phase (Supplemental Figure 4) as well as during the active repair phase (Figure 4B) following CCl_4_ administration. By *in situ* hybridization, the expression pattern of these two Wnts appeared to radiate outwards from the central vein endothelial cells and into the locally damaged tissue, but not beyond the injury border. Of note, the expression pattern of Wnt9b, normally restricted to central vein endothelial cells, is significantly broadened following injury (Figure 4B). The observation that Wnt2 and Wnt9b expression persists throughout the injury and repair phases after CCl_4_, combined with the proximity of these Wnts to the injury border, make these Wnts likely candidates as the ligands which activate the post-injury Wnt response in peri-injury Axin2+ hepatocytes.

### Endothelial cells are the major source of Wnts after injury

Our *in situ* hybridization studies (Supplemental Figure 4) suggested that non-parenchymal cells were the major source of Wnt ligands following injury. To further identify the cellular source of Wnt ligands after acute CCl_4_ injury, we conducted single cell RNA sequencing (scRNA-seq) on liver non-parenchymal cells in day 3 CCl_4_-injured mouse livers. T-distributed stochastic neighbor embedding (tSNE) analysis identified seven clusters of non-parenchymal cells in the injured liver (Figure 5A). Our analysis showed that upon injury, Wnt2 and Wnt9b are highly expressed in endothelial cells while Wnt4 is expressed in stellate cells (Figure 5B). Very few Wnt5a-expressing cells were captured in our scRNA-seq experiments (Supplemental Figure 5F).

**Figure 5.**
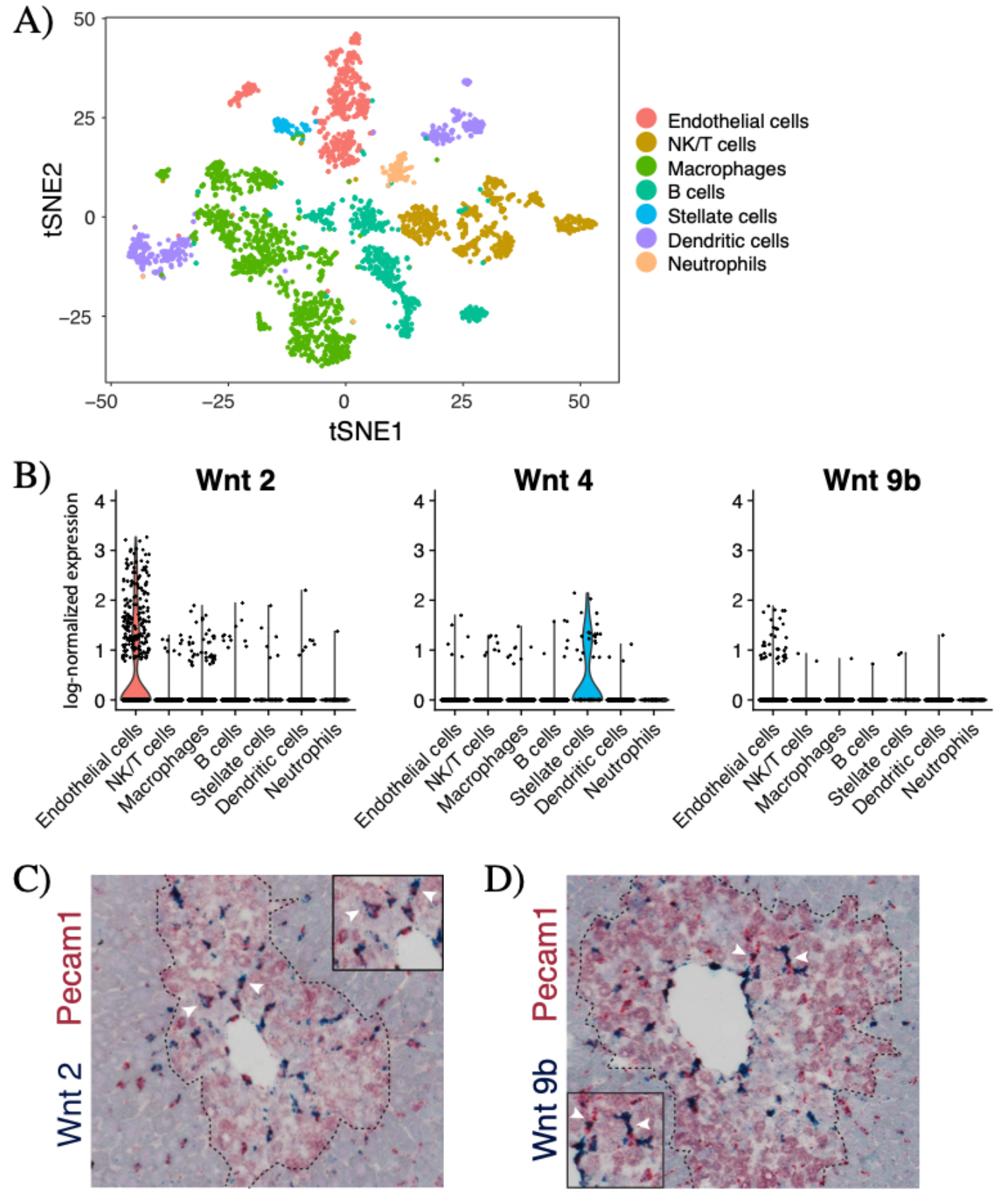
Endothelial cells are the major source of Wnt2 and Wnt9b after injury. (A) T-distributed stochastic neighbor embedding (tSNE) plot showing seven clusters of non-parenchymal cells in the day 3 CCl_4_-injured (n=3115 cells) livers. Colors denote different cell types as shown in the legend. Markers used to identify specific cell types are shown in Supplemental Figure 5D. (B) Violin plots of log-normalized expression (natural logarithm of 1+counts per 10,000) of Wnt2, Wnt4, and Wnt9b in the injured liver, categorized by cell type. Double *in situ* hybridization of Pecam1 (red) and (C) Wnt2 (blue) and (D) Wnt9b (blue) shows significant overlap between Pecam1 these two Wnts. Arrowheads denote examples of double-positive cells. Dashed line denotes injury border. N=3 animals for all studies shown.

To localize the Wnt producing cells in the injured liver lobule, we conducted double *in situ* hybridization for Wnts 2, 4, 5a, 9b and specific cell markers for endothelial cells (Pecam1), Kupffer cells (Adgre1), and stellate cells (Reln) (Supplemental Figure 6). Consistent with the scRNA-seq results, double *in situ* hybridization revealed that in the injured liver, Wnt2 and Wnt9b expression strongly overlapped with Pecam1 expression (Figure 5C & 5D), but not with Adgre1 or Reln expression (Supplemental Figure 6). A subset of Wnt4 expressing cells and a subset of Wnt5a expressing cells also expressed Reln, however neither of these two Wnts showed significant overlap with Pecam1 or Adgre1 expression (Supplementary Figure 6). Thus, after injury, endothelial cells are the major source of Wnt2 and Wnt9b while stellate cells specifically express Wnt4.

### Wnt signaling is required for injury repair

To test whether injury-induced Wnt signaling is required for tissue repair following acute CCl_4_, we interfered with intracellular Wnt signaling response by conditionally knocking out β-catenin in Wnt-responding cells in Axin2Cre^ERT2^; β-catenin^flox/Δ^ (herein Bcat-cKO) mice after CCl_4_ administration. Following tamoxifen-induced β-catenin knockout, Axin2 expression was significantly downregulated in Bcat-cKO animals after injury compared to controls (Figure 6A). Furthermore, *in situ* hybridization showed that Axin2 mRNA was specifically downregulated in peri-injury hepatocytes (Figure 6B). indicating that loss of β-catenin in Axin2+ cells effectively dampened the Wnt response in these cells. H&E staining revealed that while both control and Bcat-cKO animals exhibited similar initial injury response to 1mL/kg CCl_4_ (Supplemental Figure 8), Bcat-cKO livers appeared significantly more damaged when compared to control livers by 4 days following injury (Figure 6C & 6D). This result suggests that loss of Wnt response in peri-injury hepatocytes led to a delay in the repopulation of the damaged lesion.

**Figure 6.**
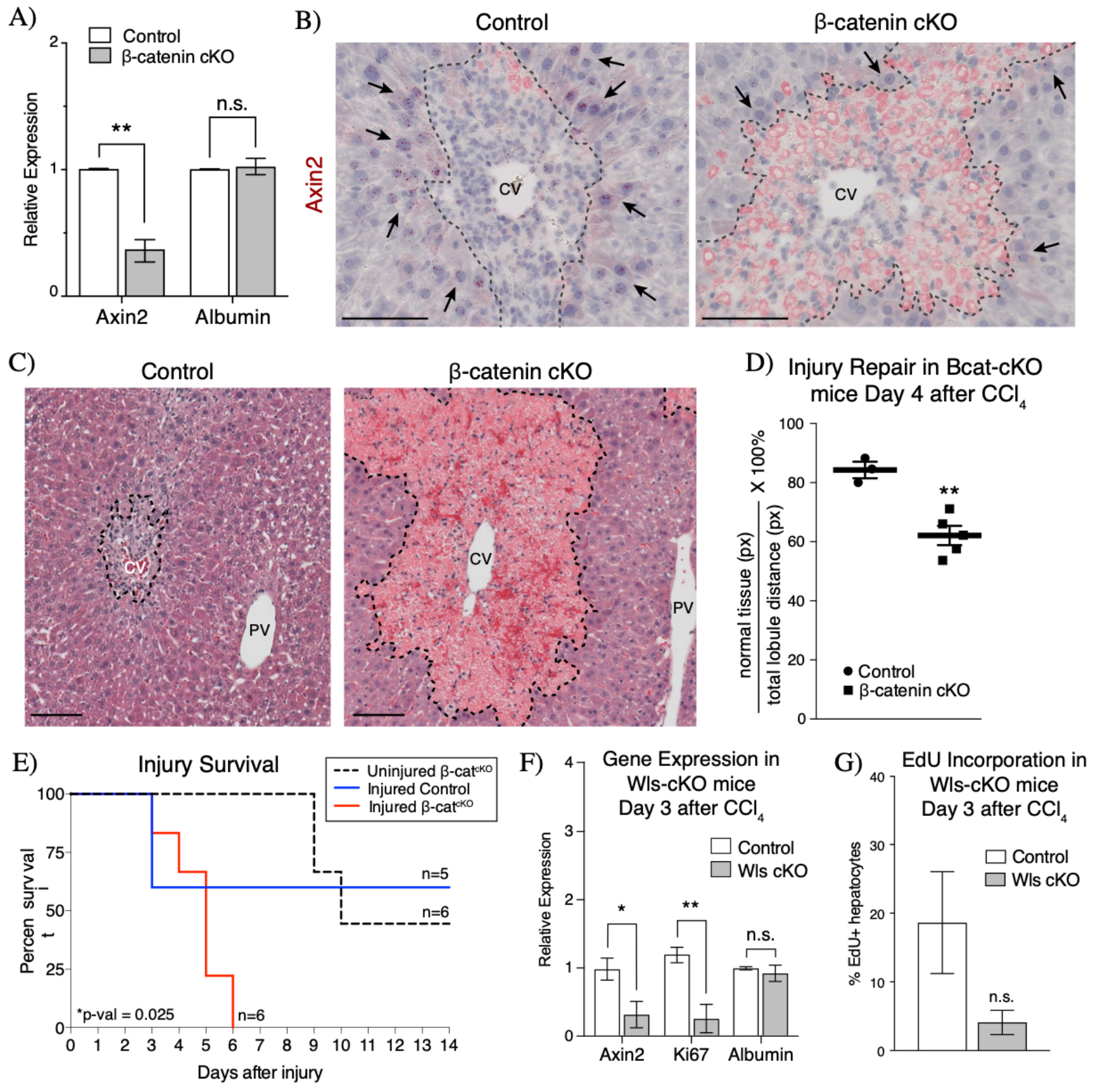
Wnt/β-catenin signaling is necessary for injury repair and survival. (A) Inducible knockout of β-catenin in the CCl_4_ injured liver results in significant decrease in Axin2 expression as measured by whole liver qRT-PCR. (B) Axin2 expression is specifically downregulated in peri-injury hepatocytes (arrows) upon loss of β-catenin in injured Bcat-cKO mouse livers (n=3 control mice, n=5 Bcat-cKO mice). (C) H&E staining morphology of injured control and Bcat-cKO livers at 4 days after injury. (D) Quantification of the extent of parenchymal repopulation based on image analysis of H&E stained sections of injured livers (see Methods) show that injured B-cat cKO livers are significantly less repaired at 4 days following injury than compared to injured control livers. (E) Loss of β-catenin ultimately results in lethality following CCl_4_ injury. (F) Loss of Wntless in endothelial cells results in decreased Axin2 and Ki67 expression by whole liver qRT-PCR at 3 days after injury. (G) A concurrent decrease in hepatocyte EdU incorporation is also observed in Wls-cKO animals (n=4 control mice, n=3 Wls cKO mice). Error bars show S.E.M. *p<0.05, **p<0.01 by two tailed t-test, n.s. = not statistically significant. 100μm scale bars. CV = central vein, PV = portal vein. Dashed line represents injury border.

To determine whether this initial delay in injury repair persists throughout the injury repair process, we conducted experiments to assess the rate of repair at later time points following CCl_4_ administration. However, we found instead that injured Bcat-cKO animals showed a higher rate of injury-induced lethality. While 3 out of 5 injured control animals survived at least 2 weeks after CCl_4_ injury, Bcat-cKO animals had a median survival of 5 days after injury (Figure 6E). It is known that loss of β-catenin from Axin2+ cells also negatively affects intestinal homeostasis, and thus it is possible that the early lethality observed in injured Bcat-cKO animals could be attributable to extra-hepatic effects such as that in the intestines. However, uninjured Bcat-cKO animals had a median survival of 10 days following β-catenin knockout, significantly higher than the median survival of injured Bcat-cKO animals (Figure 6E). Together these data suggest that loss of Wnt response in peri-injury hepatocytes following acute CCl_4_ injury results in a delay in the repair of damaged tissue, which ultimately leads to lethality from CCl_4_ toxicity.

To test further the necessity of Wnt signaling in injury repair, we examined whether endothelium-secreted Wnt ligands are critical by conditionally deleting Wntless (Wls) in Cdh5(PAC)Cre^ERT2^; Wls^flox/Δ^ (herein Wls-cKO) mice. This knockout approach was effective in blocking injury-induced Wnt activation, as shown by decreased Axin2 expression in Wls-cKO livers compared to control livers (Figure 6F). We also found that coincident with decreased Axin2 expression, Wls-cKO animals also exhibited decreased Ki67 expression levels when compared to control animals (Figure 6F). This result suggests that injury-induced cell proliferation is decreased upon loss of endothelial Wls. Consistent with the decreased Ki67 expression, hepatocyte incorporation of the thymidine-analog, EdU, was also lower at day 3 following injury in Wls-cKO animals compared to controls (Figure 6G, 4.1%±1.8% Wls-cKO vs. 18.6%±7.4% control). However, neither injured control nor injured Wls-cKO animals survived past day 3 after injury, likely due to increased sensitivity to CCl_4_ given the mixed background strain of these animals (31). Thus, while hypersensitivity to CCl_4_ in these mice prevented analysis at later injury time points, we did observe downregulation of Ki67 and decreased hepatocyte EdU incorporation in Wls-cKO animals early on after injury. These data suggest that block of endothelial Wnt secretion results in an overall decrease in the proliferative response of hepatocytes following injury and that post-injury endothelium-derived Wnt signaling is necessary for injury-induced hepatocyte proliferation.

## Discussion

Adult tissue-specific stem cells maintain organ homeostasis and are activated following injury to mediate tissue repair (32,33). However, how does tissue repair proceed upon targeted ablation of tissue-specific stem cells? In the intestines, Lgr5+ intestinal epithelial cells form a population of mitotically active intestinal stem cells that maintain intestinal homeostasis (34). However, upon ablation of these highly proliferate stem cells by irradiation, a population of otherwise quiescent Bmi1-expressing cells facilitate repair and re-epithelization (35). Thus, in the intestines, there exists a distinct “back-up” reservoir of cells that can contribute to tissue repair when the stem cell population active during homeostasis is lost.

We have previously shown that pericentral Axin2+ hepatocytes contribute to normal liver homeostasis in the uninjured mouse liver (12). Here, using the acute CCl_4_ liver injury model, we efficiently eliminated these pericentral Axin2+ hepatocytes. We found that injury results in increased expression in an expanded region of endogenous Wnt2 and Wnt9b. This injury-induced Wnt signaling activation results in a *de novo* Wnt response, in the form of Axin2 expression, in mid-lobular hepatocytes bordering the damaged local tissue. Peri-injury Axin2 expression is maintained throughout the repair time course and sustained active Wnt signaling occurs at the dynamic boundary between damaged and undamaged liver tissue after injury. Our findings show that when the homeostatic source of hepatocytes is eliminated by CCl_4_ injury, another population of Wnt-activated peri-injury hepatocytes can serve as the source of new hepatocytes for repair.

Acute CCl_4_ injury led to increased expression of Wnt2 and Wnt9b, the same Wnts that are normally expressed by centrilobular endothelial cells in the uninjured liver (12,36). We took an unbiased scRNA-seq approach to identify the cellular sources of Wnts after injury and found that endothelial cells are the primary source of Wnt2 and Wnt9b following injury. These scRNA-seq results were supported by double *in situ* experiments which showed significant overlap in the expression of these Wnts with Pecam1 expression in sinusoidal endothelial cells. Our functional studies also showed that in vivo blockade of endothelial Wnt secretion resulted in a dampening of the proliferative response following injury. Together, our Pindings are in line with previous reports that_have highlighted the importance of endothelial cells as a paracrine signaling center for regulating hepatocyte proliferation following injury (16,30,37–39). Activation of the stromal derived factor 1 (SDF1) receptor CXCR7 on liver sinusoidal endothelial cells promotes a pro-regenerative environment by inducing HGF and Wnt2 production following injury (30). On the other hand, activation of the endothelial shear-stress inducible transcription factor Krüppel-like factor 2 (KLF2) upon injury inhibits hepatocyte proliferation, in part by promoting activin A production (38). Therefore, paracrine signaling derived from liver endothelial cells serves as a check and balance system in regulating injury-induced hepatocyte proliferation.

Strikingly, we found that loss of the peri-injury hepatocyte Wnt response ultimately leads to failure to recover from CCl_4_ injury. This result is in contrast to what has been observed in the context of partial hepatectomy, where knockout of β-catenin in hepatocytes led to an initial delay in hepatocyte entry into S-phase but did not ultimately result in significant changes in liver mass recovery or survival after hepatectomy (14,40).

Our Pindings show that following a *localized* injury (i.e. CCl_4_), only select hepatocytes, specifically those closest to the damaged tissue, are more likely to become activated and undergo proliferation, while hepatocytes farther from the injury site remain quiescent. This is in contrast to the pan-hepatocyte proliferative response observed after a global liver injury such as partial hepatectomy. Additionally, our results show that most hepatocytes have the *capacity* to respond to a Wnt signal. This inherent capacity for hepatocellular response to Wnt signaling is biologically significant given the liver’s increased risk for injury as a central detoxification organ and the major role of Wnt/β-catenin signaling as a regulator for liver tissue repair.

## Acknowledgements

We thank T. Desai, J. Sage, L. Xing, S. Tan, H. Takase, J. Tsai, T. Anbarchian, E. Rim, E. Rulifson, W.C. Peng, and K. Loh for helpful discussions on experimental design and data analysis; G. Karnam, D. Burhan and the UCSF Liver Center for assistance with liver cell isolations; P. Lovelace for assistance with Plow cytometry, and K. Shaw for assistance in reagent and supplies acquisition.

## Supplemental Figures

**Supplemental Figure 1.**
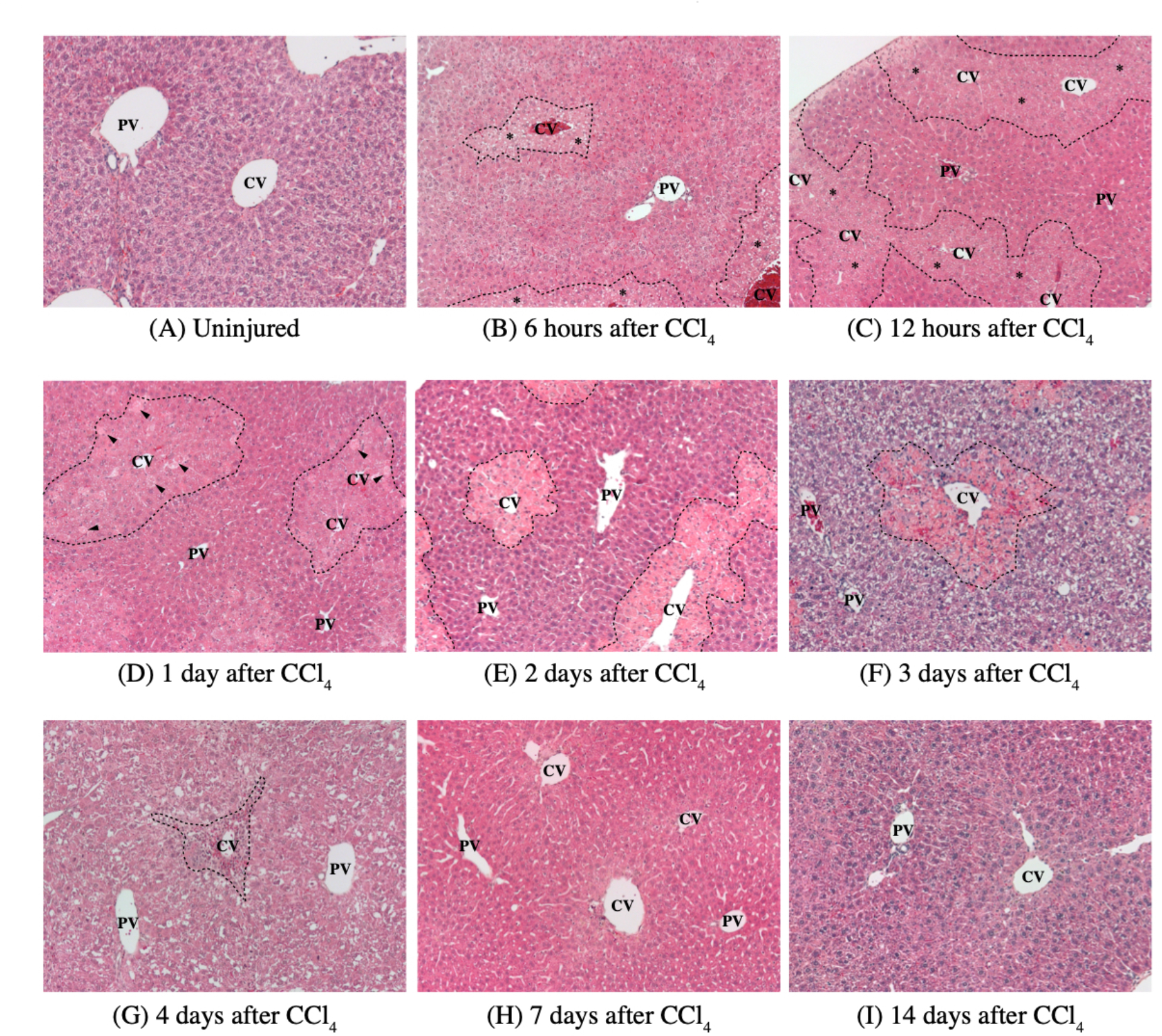
Hematoxylin and eosin stained morphology timeline of CCl_4_ induced liver injury. (A) H&E of the uninjured adult mouse liver. (B) 6 hours following CCl_4_ administration, hepatocytes closest to the central vein (inside dashed lines) begin to show signs of steatosis (*). (C) By 12 hours, the affected area broadens and more hepatocytes show signs of steatosis (*). (D) 1 day after CCl_4_ administration, apoptotic cells (arrowheads) are observed within the affected pericentral region. (E) Massive centrilobular necrosis becomes obvious by day 2 after CCl_4_ administration (inside dashed lines). (F) The repair phase begins around 3 days following CCl_4_ administration. Inflammatory infiltration is readily visible, as macrophages and neutrophils invade the necrotic debris. (G) By day 4, the injured area shrinks in size due to the proliferation of hepatocytes and the clearing out of the necrotic debris. (H) By 1 week after CCl_4_, the liver lobule looks most normal. Remnant inflammatory infiltration can be observed at some central veins. (I) The normal histology of the liver lobule is restored by 2 weeks after CCl_4_. CV = central vein, PV = portal vein. All images taken with a 10x objective.

**Supplemental Figure 2.**
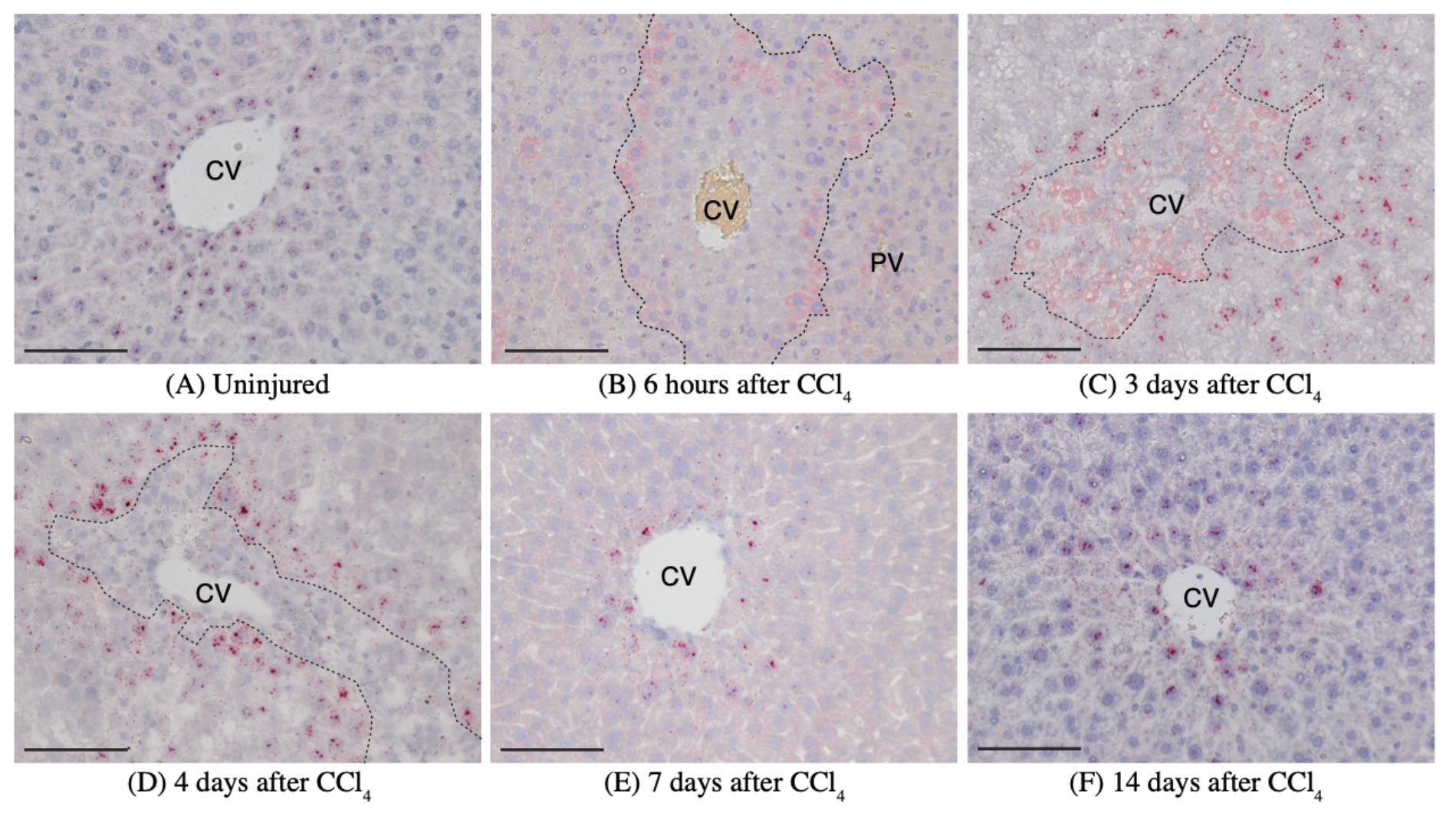
Axin2 mRNA *in situ* hybridization timeline of CCl_4_ induced liver injury. (A) Axin2 expression (red dots) is limited to pericentral hepatocytes in the uninjured liver. (B) Loss of Axin2 expression in the pericentral region is evident by 6 hours after CCl_4_. (C & D) Robust Axin2 expression is observed in mid-lobular hepatocytes adjacent to the injury border by day 3. (E & F) Normal Axin2 expression pattern in pericentral hepatocytes is re-established upon injury recovery. CV = central vein, PV = portal vein. Dashed lines denote injury border. 100μm scale bars.

**Supplemental Figure 3.**
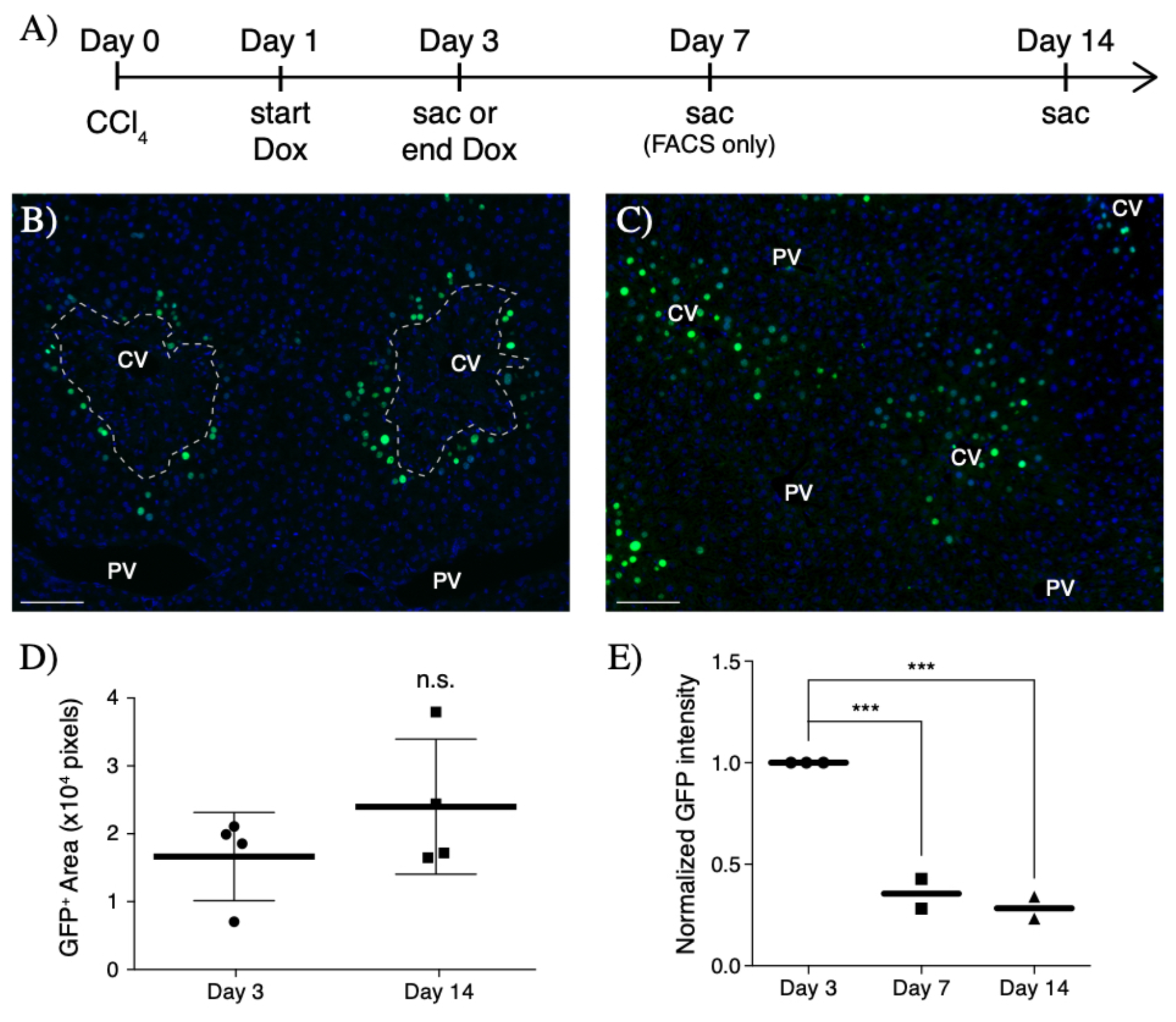
Label dilution studies in injured Axin2-rtTA; Tet-O-H2BGFP reporter mice. (A) Schematic of CCl_4_ injury and doxycycline (Dox) “pulse-chase” scheduling. (B) Peri-injury Axin2+ hepatocytes (green) are labeled after 48 hours of 1 mg/mL doxycycline chase following CCl_4_ (day 3). (C) After an 11 day “chase” period, newly repopulated pericentral region contains predominantly H2BGFP-labeled hepatocytes. At this timpeoint, it is evident that H2BGFP signal intensity is significantly lower than that observed at day 3, however GFP signal for the image shown was enhanced in order for dim H2BGFP signal to be visualized. 100um scale bars shown. CV = central vein, PV = portal vein. Dashed lines represent injury border. (D) There was an overall increase in the amount of labeled cells, as measured by GFP+ nuclear area, over the injury repair course, though this increase did not reach statistical significance (n=4 animals analyzed per timepoint, error bars represent S.E.M.). (E) Quantification of GFP intensity by Vlow cytometry. GFP mean fluorescence intensity (MFI) was normalized to that of the GFP+ population observed in day 3 samples. Each symbol represents the MFI for one animal, n=3 animals for day 3, n=2 for days 7, and n=2 for day 14. ***p-val<0.001 by multiple comparisons using one-way ANOVA.

**Supplemental Figure 4.**
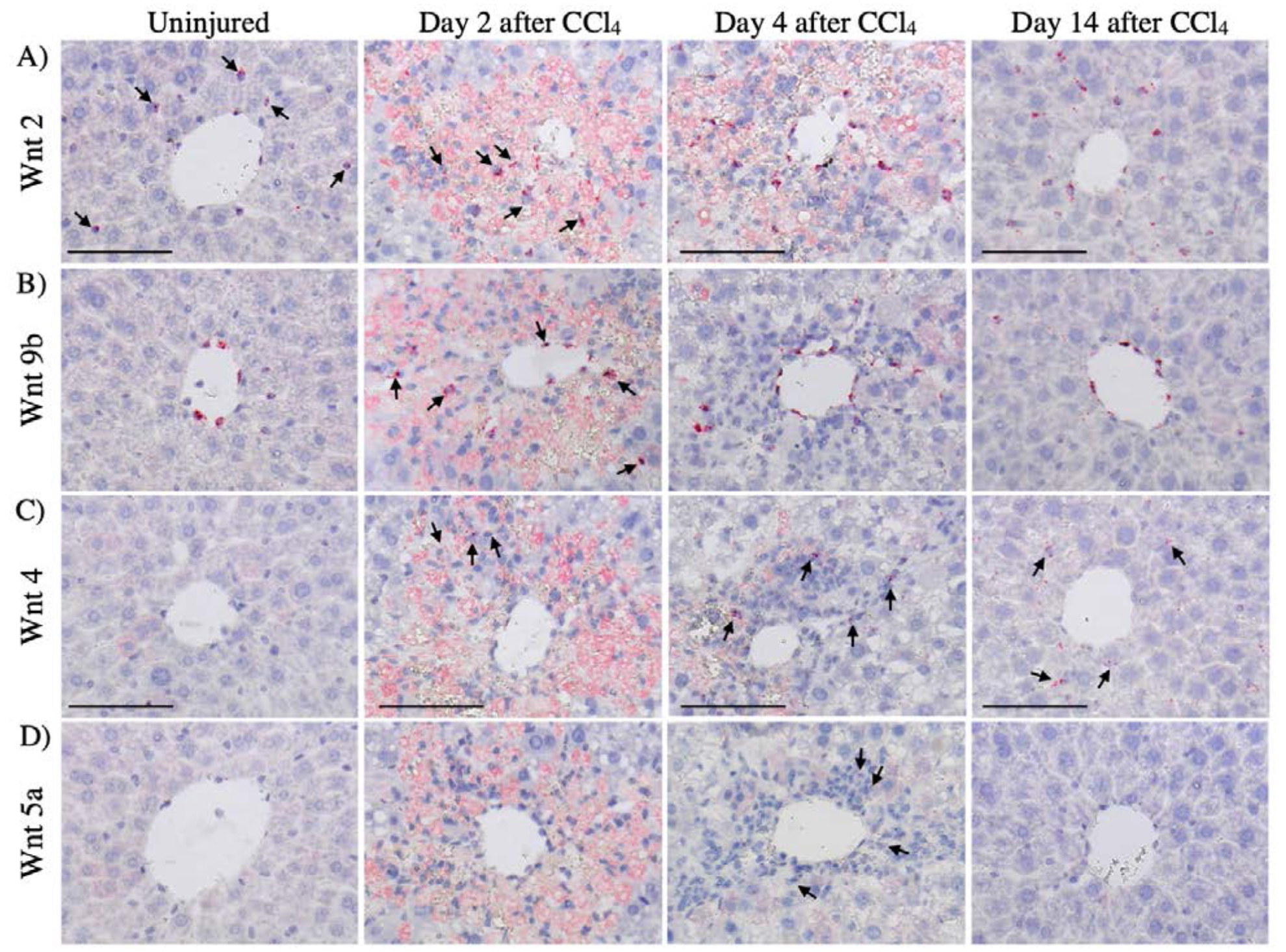
mRNA *in situ* hybridization for Wnt2, Wnt9b, Wnt4, and Wnt5a. (A) Wnt2 is expressed in the injured tissue as well as in the uninjured and recovered liver. (B) Wnt9b expression broadens during repair. (C) Wnt4 is expressed in the injured area when inflammatory infiltration is at its peak (day 4 after CCl_4_). (D) Wnt5a is also expressed in the injured area only when inflammatory infiltration is at its peak (day 4 after CCl_4_). Diffuse pink staining found in the day 3 column is non-speciflc stain trapping by damaged tissue. Positive signal are the red puncta. 100μm scale bars. Arrows indicate examples of positive *in situ* signal.

**Supplemental Figure 5.**
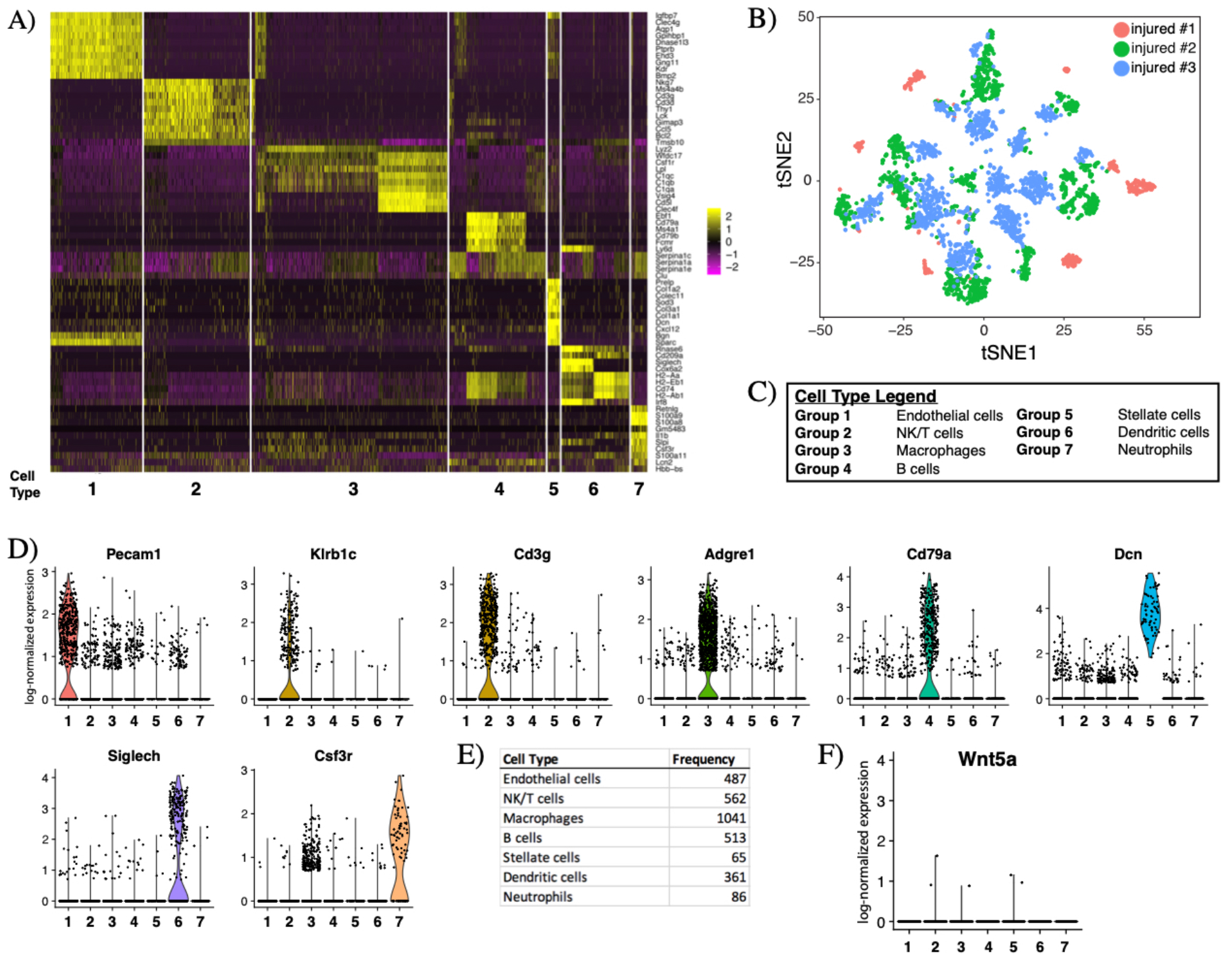
(A) Single cell RNA-seq derived gene expression profile of non-parenchymal cells in the day 3 CCl_4_-injured liver; each row represents an individual gene, each column represents an individual cell. Yellow denotes more expression, magenta denotes less expression. Cells are grouped by cell type as shown in the (C) legend. (B) tSNE plot showing contribution from individual animals, color coded according to leged in upper right corner of the plot. (D) Violin plots of log-normalized expression (natural logarithm of 1+counts per 10,000) of cell type specific markers used to identify seven distinct non-parenchymal cell clusters after injury. (E) Frequency of specific cell types sampled in scRNA-seq analysis. (F) Violin plots of log-normalized expression of Wnt5a show that very few Wnt5a expressing cells were captured in our scRNA-seq experiments.

**Supplemental Figure 6.**
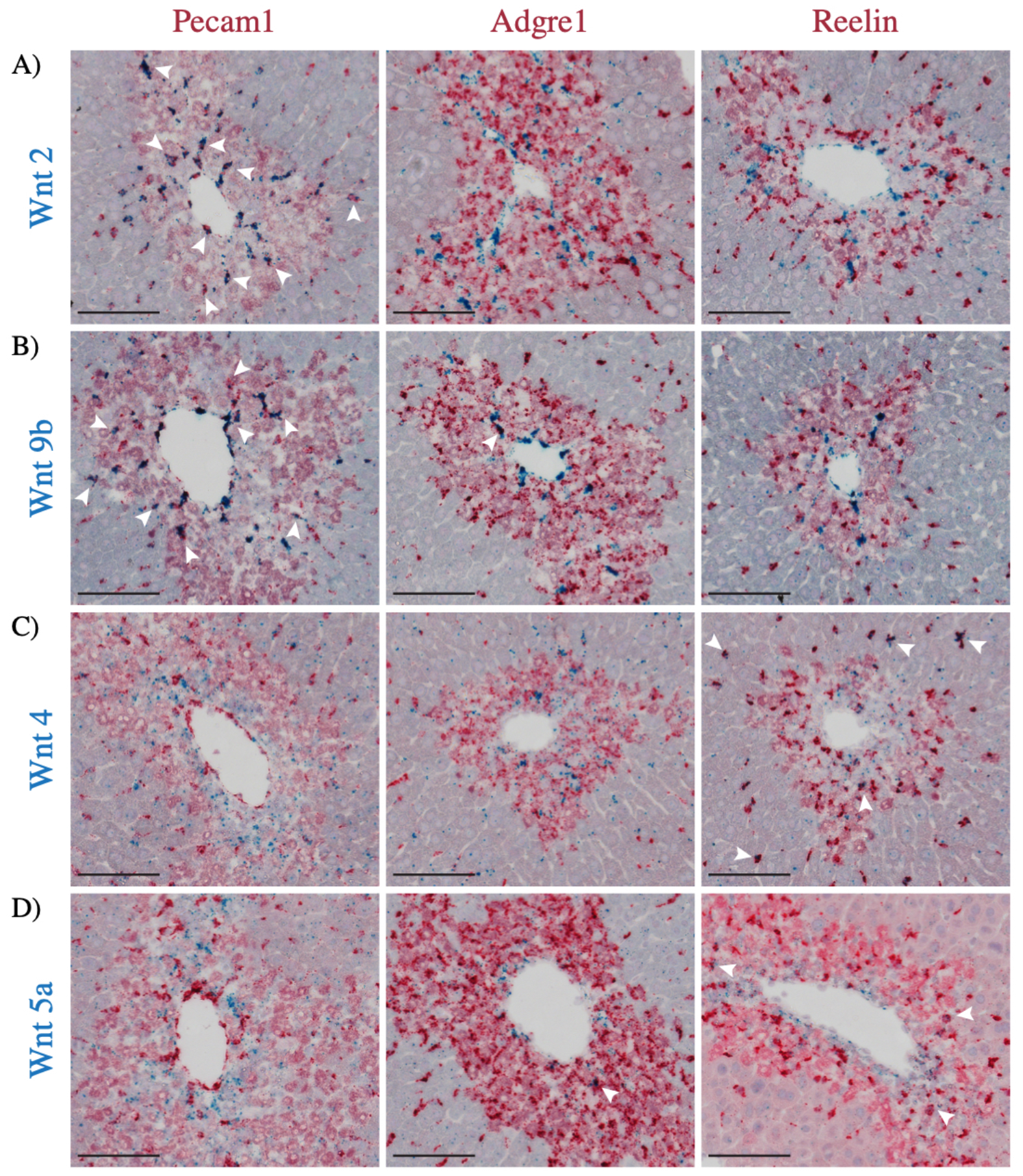
Double mRNA *in situ* hybridization for Wnt 2, 4, 5a, 9b and Pecam1 (endothelial cells), Adgre1 (Kupffer cells/macrophages), or Reelin (stellate cells) in the day 3 injured liver. (A) Wnt2 is expressed by Pecam1+ cells but not by Adgre1 or Reln positive cells. (B) Wnt9b is expressed by Pecam1+ cells but not by Reln positive cells. There is 1 cell that expresses both Wnt9b and Adgre1 in the image shown. (C) Wnt4 is expressed by Reln+ cells, but not by Adgre1 or Pecam1 positive cells. (D) Wnt5a is expressed by Reln+ cells, but not by Pecam1 positive cells. There is 1 cell that is double positive for Wnt5a and Adgre1 in the image shown. Blue puncta represents positive Wnt signal, red puncta represents positive cell-type specific marker signal. Diffuse pink staining found in the pericentral necrotic tissue is due to color trapping by damaged tissue. 100μm scale bars. Arrowheads indicate examples of double-positive *in situ* signal.

**Supplemental Figure 7.**
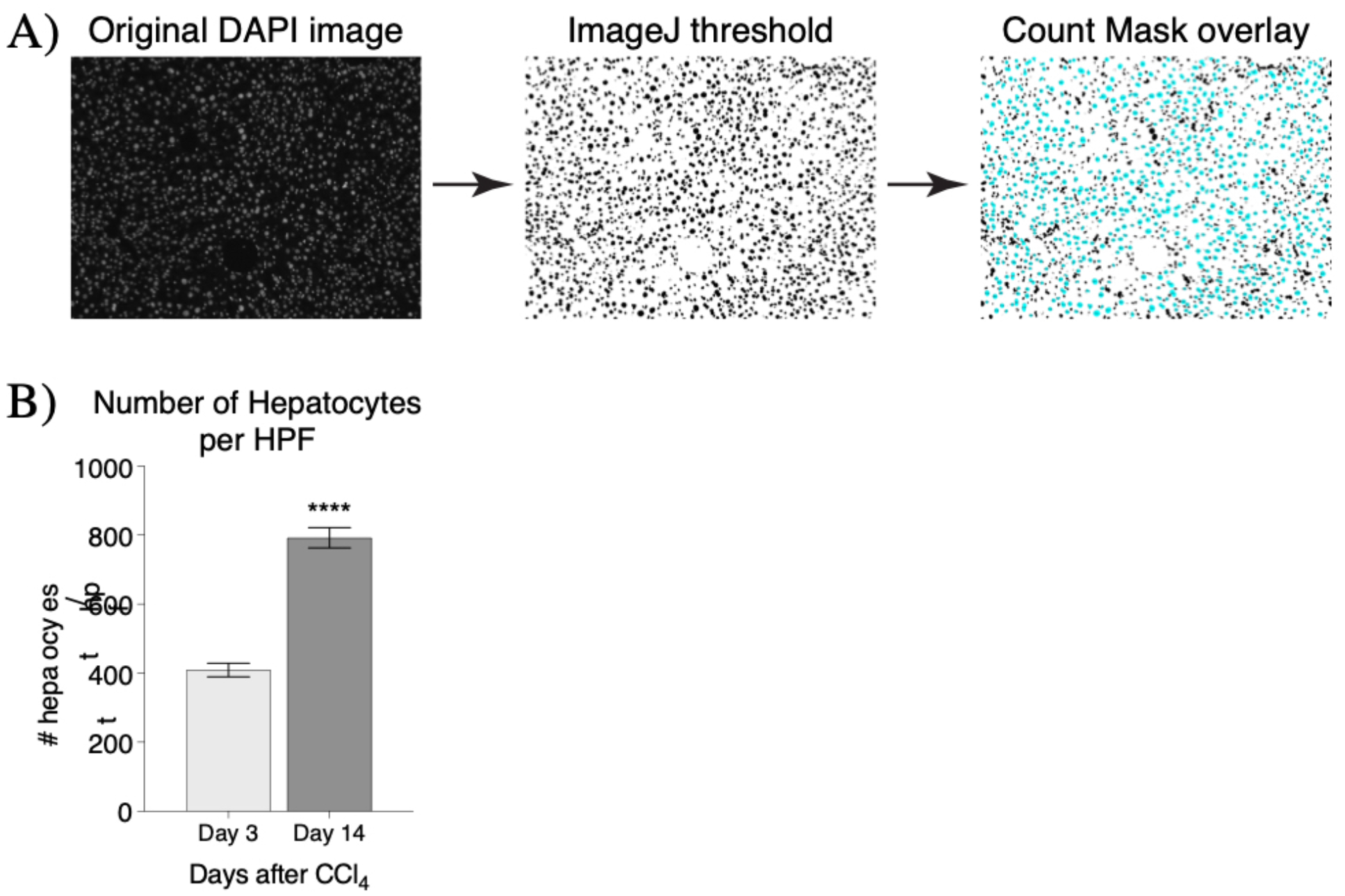
Hepatocyte number following injury and upon injury repair. (A) Diagram of ImageJ automated thresholding and particle analysis workflow that was used to generate hepatocyte count data. (B) The number of hepatocytes significantly increased between 3 days (n=3) and 14 days (n=4) following CCl_4_ injury. Error bars show S.E.M. ****p<0.0001 by two-tailed t-test.

**Supplemental Figure 8.**
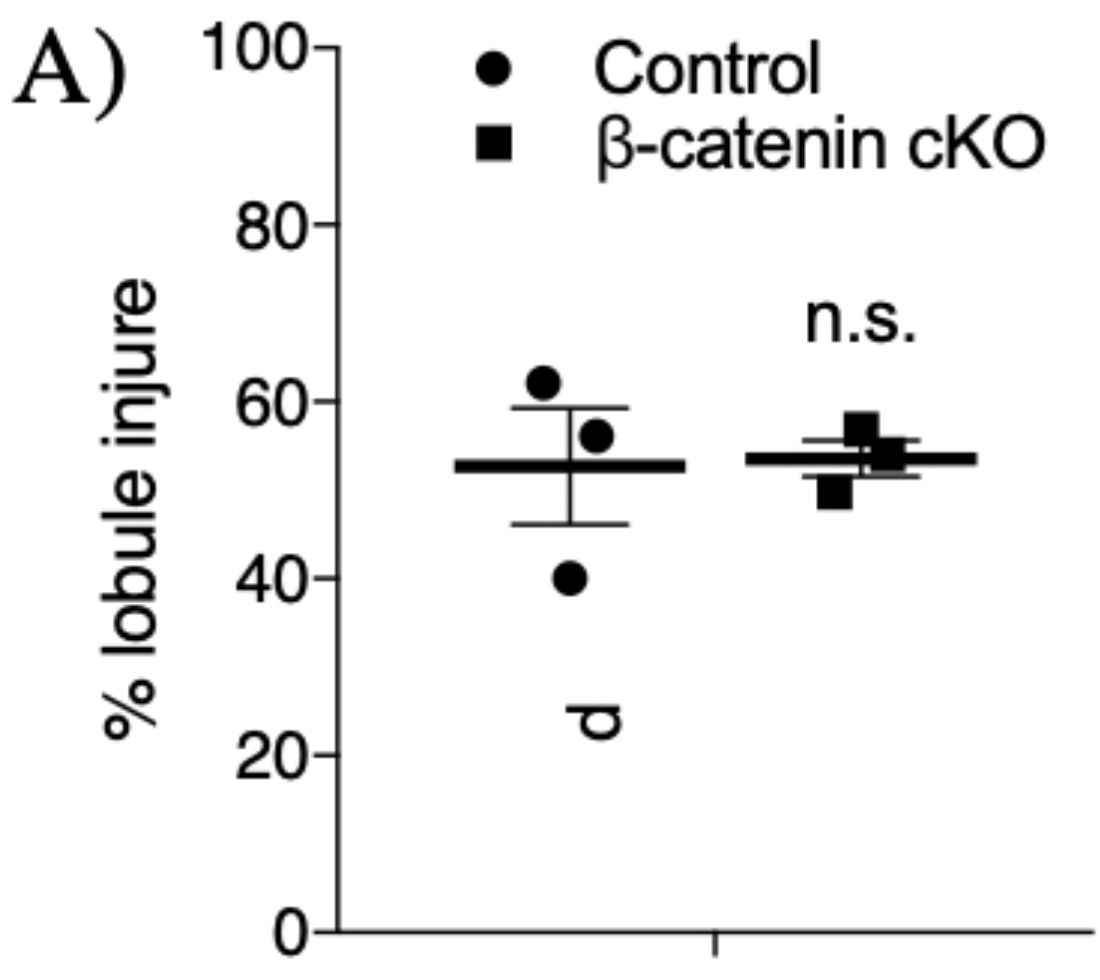
Initial injury response to CCl_4_ in control and Bcat-cKO mice. (A) Control and Bcat-cKO mice exhibited similar extent of parenchymal injury following 1mL/ kg CCl_4_ administration on injury day 2. Error bars show S.E.M., n.s. = not statistically significant (p=0.9) by two-tailed t-test for n=3 control and n=3 Bcat-cKO animals.

## References

1. Malato Y, Naqvi S, Schürmann N, Ng R, Wang B, Zape J, et al. Fate tracing of mature hepatocytes in mouse liver homeostasis and regeneration. J. Clin. Invest. 2011;121:4850–4860.

2. Yanger K, Knigin D, Zong Y, Maggs L, Gu G, Akiyama H, et al. Adult Hepatocytes Are Generatedby Self-DuplicationRather than Stem Cell Differentiation. Stem Cell. 2014;:1–22.

3. Newsome PN, Hussain MA, Theise ND. Hepatic oval cells: helping redefine a paradigm in stem cell biology. Curr. Top. Dev. Biol. 2004;61:1–28.

4. Huch M, Dorrell C, Boj SF, Van Es JH, Li VSW, van de Wetering M, et al. In vitro expansion of single Lgr5+ liver stem cells induced by Wnt-driven regeneration. Nature. 2013;494:247–250.

5. Tanimizu N, Mitaka T. Re-evaluation of liver stem/progenitor cells. Organogenesis. 2014;10:208–215.

6. Tarlow BD, Finegold MJ, Grompe M. Clonal tracing of Sox9+ liver progenitors in mouse oval cell injury. Hepatology. 2014;60:278–289.

7. Raven A, Lu W-Y, Man TY, Ferreira-Gonzalez S, O’Duibhir E, Dwyer BJ, et al. Cholangiocytes act as facultative liver stem cells during impaired hepatocyte regeneration. Nature. 2017;547:350–354.

8. Micsenyi A, Tan X, Sneddon T, Luo J-H, Michalopoulos GK, Monga SPS. Beta-catenin is temporally regulated during normal liver development. Gastroenterology. 2004;126:1134–1146.

9. Apte U, Zeng G, Thompson MD, Muller P, Micsenyi A, Cieply B, et al. beta-Catenin is critical for early postnatal liver growth. Am. J. Physiol. Gastrointest. Liver Physiol. 2007;292:G1578–85.

10. Thompson MD, Monga SPS. WNT/β-catenin signaling in liver health and disease. Hepatology. 2007;45:1298–1305.

11. Tan X, Yuan Y, Zeng G, Apte U, Thompson MD, Cieply B, et al. Beta-catenin deletion in hepatoblasts disrupts hepatic morphogenesis and survival during mouse development. Hepatology. 2008;47:1667–1679.

12. Wang B, Zhao L, Fish M, Logan CY, Nusse R. Self-renewing diploid Axin2(+) cells fuel homeostatic renewal of the liver. Nature. 2015;524:180–185.

13. Yang J, Mowry LE, Nejak-Bowen KN, Okabe H, Diegel CR, Lang RA, et al. Beta-catenin signaling in murine liver zonation and regeneration: A Wnt-Wnt situation! Hepatology. 2014;60:964–976.

14. Sekine S, Gutiérrez PJA, Yu-Ang Lan B, Feng S, Hebrok M. Liver-specific loss of β-catenin results in delayed hepatocyte proliferation after partial hepatectomy. Hepatology. 2007;45:361–368.

15. Nejak-Bowen KN, Thompson MD, Singh S, Bowen WC, Dar MJ, Khillan J, et al. Accelerated liver regeneration and hepatocarcinogenesis in mice overexpressing serine-45 mutant beta-catenin. Hepatology. 2010;51:1603–1613.

16. Ding B-S, Nolan DJ, Butler JM, James D, Babazadeh AO, Rosenwaks Z, et al. Inductive angiocrine signals from sinusoidal endothelium are required for liver regeneration. Nature. 2010;468:310–315.

17. Planas-Paz L, Orsini V, Boulter L, Calabrese D, Pikiolek M, Nigsch F, et al. The RSPO-LGR4/5-ZNRF3/RNF43 module controls liver zonation and size. Nat. Cell Biol. 2016;18:467–479.

18. Rocha AS, Vidal V, Mertz M, Kendall TJ, Charlet A, Okamoto H, et al. The Angiocrine Factor Rspondin3 Is a Key Determinant of Liver Zonation. CellReports. 2015;13:1757–1764.

19. Lustig B, Jerchow B, Sachs M, Weiler S, Pietsch T, Karsten U, et al. Negative Feedback Loop of Wnt Signaling through Upregulation of Conductin/Axin2 in Colorectal and Liver Tumors. Molecular and Cellular Biology. 2002;22:1184–1193.

20. van Amerongen R, Bowman AN, Nusse R. Developmental stage and time dictate the fate of Wnt/β-catenin-responsive stem cells in the mammary gland. Cell Stem Cell. 2012;11:387–400.

21. Madisen L, Zwingman TA, Sunkin SM, Oh SW, Zariwala HA, Gu H, et al. A robust and high-throughput Cre reporting and characterization system for the whole mouse brain. Nat. Neurosci. 2010;13:133–140.

22. Brault V, Moore R, Kutsch S, Ishibashi M, Rowitch DH, McMahon AP, et al. Inactivation of the beta-catenin gene by Wnt1-Cre-mediated deletion results in dramatic brain malformation and failure of craniofacial development. Development. 2001;128:1253–1264.

23. Carpenter AC, Rao S, Wells JM, Campbell K, Lang RA. Generation of mice with a conditional null allele for Wntless. Genesis. 2010;48:554–558.

24. F Schwenk Ubkr. A cre-transgenic mouse strain for the ubiquitous deletion of loxP-flanked gene segments including deletion in germ cells. Nucleic Acids Research. 1995;23:5080.

25. Wang Y, Nakayama M, Pitulescu ME, Schmidt TS, Bochenek ML, Sakakibara A, et al. Ephrin-B2 controls VEGF-induced angiogenesis and lymphangiogenesis. Nature. 2010;465:483–486.

26. Livak KJ, Schmittgen TD. Analysis of relative gene expression data using real-time quantitative PCR and the 2(-Delta Delta C(T)) Method. Methods. 2001;25:402–408.

27. Tabula Muris Consortium, Overall coordination, Logistical coordination, Organ collection and processing, Library preparation and sequencing, Computational data analysis, et al. Single-cell transcriptomics of 20 mouse organs creates a Tabula Muris. Nature. 2018;562:367–372.

28. Ghafoory S, Breitkopf-Heinlein K, Li Q, Scholl C, Dooley S, Wölfl S. Zonation of nitrogen and glucose metabolism gene expression upon acute liver damage in mouse. PLoS ONE. 2013;8:e78262.

29. Jho E-H, Zhang T, Domon C, Joo C-K, Freund J-N, Costantini F. Wnt/beta-catenin/Tcf signaling induces the transcription of Axin2, a negative regulator of the signaling pathway. Molecular and Cellular Biology. 2002;22:1172–1183.

30. Ding B-S, Cao Z, Lis R, Nolan DJ, Guo P, Simons M, et al. Divergent angiocrine signals from vascular niche balance liver regeneration and fibrosis. Nature. 2014;505:97–102.

31. Bhathal PS, Rose NR, Mackay IR, Whittingham S. Strain differences in mice in carbon tetrachloride-induced liver injury. Br J Exp Pathol. 1983;64:524–533.

32. Relaix F, Zammit PS. Satellite cells are essential for skeletal muscle regeneration: the cell on the edge returns centre stage. Development. 2012;139:2845–2856.

33. Gregorieff A, Liu Y, Inanlou MR, Khomchuk Y, Wrana JL. Yap-dependent reprogramming of Lgr5+ stem cells drives intestinal regeneration and cancer. Nature. 2015;526:715–718.

34. Barker N, Van Es JH, Kuipers J, Kujala P, van den Born M, Cozijnsen M, et al. Identification of stem cells in small intestine and colon by marker gene Lgr5. Nature. 2007;449:1003–1007.

35. Yan KS, Chia LA, Li X, Ootani A, Su J, Lee JY, et al. The intestinal stem cell markers Bmi1 and Lgr5 identify two functionally distinct populations. Proceedings of the National Academy of Sciences. 2012;109:466–471.

36. Halpern KB, Shenhav R, Massalha H, Tóth B, Egozi A, Massasa EE, et al. Paired-cell sequencing enables spatial gene expression mapping of liver endothelial cells. Nat Biotechnol. 2018;5:279–15.

37. Hu J, Srivastava K, Wieland M, Runge A, Mogler C, Besemefelder E, et al. Endothelial Cell-Derived Angiopoietin-2Controls Liver Regeneration as aSpatiotemporal Rheostat. Science. 2014;343:416–419.

38. Manavski Y, Abel T, Hu J, Kleinlützum D, Buchholz CJ, Belz C, et al. Endothelial transcription factor KLF2 negatively regulates liver regeneration via induction of activin A. Proceedings of the National Academy of Sciences. 2017;114:3993–3998.

39. Gomez-Salinero JM, Rafii S. Endothelial cell adaptation in regeneration. Science. 2018;362:1116–1117.

40. Tan X, Behari J, Cieply B, Michalopoulos GK, Monga SPS. Conditional Deletion of β-Catenin Reveals Its Role in Liver Growth and Regeneration. Gastroenterology. 2006;131:1561–1572.

